# Phenotypic dominance emerges from activity-fitness functions and molecular interactions

**DOI:** 10.1101/2025.03.21.644634

**Authors:** Tushar Pal, Soham Dibyachintan, Angel F. Cisneros, Christian R. Landry

**Affiliations:** Département de biochimie, de microbiologie et de bio-informatique, Faculté des sciences et de génie, Université Laval, G1V 0A6, Québec, Canada; Institut de biologie intégrative et des systèmes, Université Laval, G1V 0A6, Québec, Canada; PROTEO, Le regroupement québécois de recherche sur la fonction, l’ingénierie et les applications des protéines, Université Laval, G1V 0A6, Québec, Canada; Centre de recherche sur les données massives, Université Laval, G1V 0A6, Québec, Canada; Department of Biological Sciences, Indian Institute of Science Education and Research Kolkata, Mohanpur, Nadia, West Bengal, 741246, India; Centre de recherche en infectiologie, Université Laval, G1V 0A6, Québec, Canada

**Author notes:** To whom correspondence should be addressed, Correspondance; E-mail: Angel F. Cisneros, Christian R. Landry.

**Keywords:** Phenotypic dominance, molecular dominance, mutational effects, fitness, protein interactions, homomeric proteins, fitness functions

## Abstract

Mendelian genetics provides us with a framework for studying allelic dominance relationships at a locus at the phenotypic level. These dominance relationships result from different molecular factors, including the mapping of molecular activity onto fitness. For homomeric proteins, physical interactions between alleles provide a mechanism by which one allele can have a dominant effect on the activity of the other. Here, we refer to the effect of these interactions as molecular dominance and examine how they determine total protein activity and contribute to phenotypic dominance. The relative impact of such molecular dominance effects depends on the proportion of subunits that heteromerize relative to those that form homomers. In turn, we show how the effect of physical interactions on phenotypic dominance depends on the function linking protein activity to fitness. Our results show the complex relationships between molecular and phenotypic dominance and highlight the fundamental difference in dominance landscapes for monomeric and homomeric proteins.

## Introduction

A major question in genetics is how alleles combine to produce phenotypes. Mendel’s pioneering experiments on inheritance showed that in diploid organisms, some alleles are dominant over others (Mendel 1865), the dominant allele being the one that shows the same phenotype in the homo- and heterozygotes. Dominance relationships between alleles have major consequences for human health and evolution. For example, many human disease-causing alleles are autosomal dominant (Zschocke, Byers, and Wilkie 2023; Hamosh et al. 2005), which increases the probability of phenotypic transmission of diseases between parent and offspring. In evolution, dominant deleterious or adaptive alleles are always visible to natural or artificial selection, which can accelerate their rate of loss or fixation (Fisher, Vignogna, and Lang 2021; Aggeli et al. 2022; Bautista et al. 2024; Le Poul et al. 2014). While the phenotypic consequences of dominance and recessivity are well known, the underlying molecular bases are far from being completely understood, particularly when the products of two allele copies can functionally interact.

When fitness is the trait of interest, dominance can be calculated by comparing the reproductive success (fitness) of homo- and heterozygotes (Greenberg and Crow 1960; Le Poul et al. 2014; Huber et al. 2018). The fitness of mutant homozygous and heterozygous individuals is compared to a reference homozygote, typically the wild-type (WT). By definition, the fitness of the WT homozygote (which we refer to as *ω*_aa_) is set to 1. A selection coefficient (*s*) is estimated by comparing the fitness of mutant homozygous and WT individuals, which is positive for deleterious genotypes and negative for beneficial ones. Then, a dominance coefficient (*h*, also called heterozygous effect) is inferred based on the fitness of heterozygous individuals and the selection coefficients of the corresponding homozygotes (Greenberg and Crow 1960). Alleles are typically considered recessive if 0 ≤ *h* < 0.5 and dominant if 0.5 < *h* ≤ 1. For example, in the case of a heterozygote individual harboring a mutant and a WT allele, the selection and dominance coefficients would be given by equations 1 and 2:

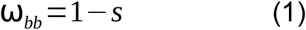

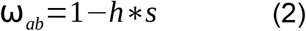

where *ω*_bb_ and *ω*_ab_ respectively indicate the fitness of the mutant homozygote and the heterozygote, *s* indicates the selection coefficient of the mutant homozygote, and *h* indicates the dominance coefficient of the mutant allele.

Different models have aimed to explain the phenotypic dominance of specific alleles. Since the contribution of a gene to a phenotype is often linked to the function (activity) and abundance of its encoded protein, these models generally posit that fitness depends on the activity summed over the two alleles. Thus, these models consider fitness functions that establish quantitative relationships between fitness and the total protein activity. Dominance can emerge directly from nonlinearities in these fitness functions (Wright 1934; Kacser and Burns 1981; Hurst and Randerson 2000; Huber et al. 2018; Di and Lohmueller 2024; Billiard, Castric, and Llaurens 2021; Manna, Martin, and Lenormand 2011; Brown et al. 2009; Xie et al. 2023). For example, consider a loss-of-function allele for an enzyme. Due to the lack of protein activity, the mutant homozygote has a very deleterious phenotype. As these models of dominance assume that the two copies are independent, a heterozygous loss-of-function mutation is expected to cause a 50% reduction in activity. However, if the 50% reduction in activity of the heterozygote reduces fitness by less than 50%, the loss-of-function allele would be recessive. Fitness functions with diminishing returns for higher activity, such as those observed in metabolic flux analysis (Hartl, Dykhuizen, and Dean 1985; Dykhuizen, Dean, and Hartl 1987), typically produce such recessive alleles. On the other hand, if the 50% reduction in activity reduces fitness by more than 50%, the loss-of-function allele would be dominant. Such outcomes are typically observed for intervals of fitness functions where higher activity provides increasing benefits, as shown for beta-lactamases that degrade antibiotics (Zimmermann and Rosselet 1977; Nikaido and Normark 1987; Lakaye et al. 1999). The functions mapping activity to fitness are thus critical for our understanding of dominance, especially since they can change for different proteins or environmental conditions (Després et al. 2022; Keren et al. 2016; Z. Wu et al. 2022; Otto et al. 2024; Qi et al. 2013).

Physical interactions between proteins encoded by alleles of the same gene are another factor that can shape the dominance effects of alleles. Approximately 20-45% of proteins self-assemble into homomers (Schweke et al. 2024), making dimerization of allelic products a common phenomenon. In diploids, the two corresponding alleles of homomeric proteins usually assemble into a mix of homo- and heteromers (Pereira-Leal et al. 2007; Walker et al. 1997) unless specific mechanisms mediate allele-specific assembly (Badonyi and Marsh 2023). These heteromers then provide a mechanism by which a mutant allele can interfere with the function of a WT copy, providing an opportunity for dominance to occur at the molecular level as well. Well-known examples of molecular dominance include dominant negative alleles. These alleles are inactive themselves and sequester WT subunits in catalytically inactive heterocomplexes (Herskowitz 1987; Reiner A. Veitia 2007; R. A. Veitia, Caburet, and Birchler 2018). Since these mutations often result in drastic losses of protein function, they have been associated with a variety of human diseases (Zschocke, Byers, and Wilkie 2023; Bergendahl et al. 2019; Backwell and Marsh 2022; Meyer-Schuman et al. 2023).

The above observations show that both fitness functions and molecular interactions contribute to the emergence of phenotypic dominance. Since current frameworks do not model physical interactions between alleles (Wright 1934; Kacser and Burns 1981; Hurst and Randerson 2000; Huber et al. 2018; Manna, Martin, and Lenormand 2011; Xie et al. 2023), it is difficult to disentangle the relative contributions of fitness functions and physical interactions to dominance. Here, we investigate how molecular dominance between alleles translates into phenotypic dominance in the context of a fitness function. We build a model that introduces molecular terms representing the reduction in the specific activity of a mutant protein relative to the WT (*r*) and the dominance of such effects on the activity of the heterodimer (*g*). These parameters are analogous to the parameters that describe the effect of mutations on organismal phenotype: the selection (*s*) and dominance (*h*) coefficients. By comparing phenotypes emerging from the activity of monomeric and homodimeric proteins, we show that molecular dominance in homodimeric proteins produces combinations of changes in activity and fitness effects that are not equally accessible to monomeric proteins. Similarly, we demonstrate that dominant mutational effects are potentiated if mutations not only impact protein activity but also make the fraction of heteromers increase relative to that of homomers. Finally, we show how the effect of molecular dominance combines with fitness functions to produce phenotypic dominance.

## Model

We built a model assuming a diploid organism, with a gene of interest, “a”, with three genotypes: 1) WT homozygous (with both alleles being the wild type), 2) mutant homozygous (with both alleles having the same mutation), and 3) heterozygous (with one allele being WT and the other being a mutant allele). From here onwards, we will refer to WT alleles as “a” and mutant alleles as “b”. Proteins “A” and “B” are synthesized from alleles “a” and “b”, respectively. Here, we will focus on the effects of physical interactions on the sum of the activity of the two proteins synthesized from the two alleles and refer to it as total protein activity (ϕ). In the case of). In the case of monomers (**Fig. 1a**), the two proteins are independent of one another. We define *r* as the reduction in activity of the mutant protein compared to the WT monomer (with specific activity of 1), as shown in equation 3:

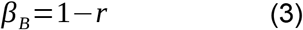

where *β*_*B*_ indicates the specific activity of the mutant monomer.

**Figure 1.**
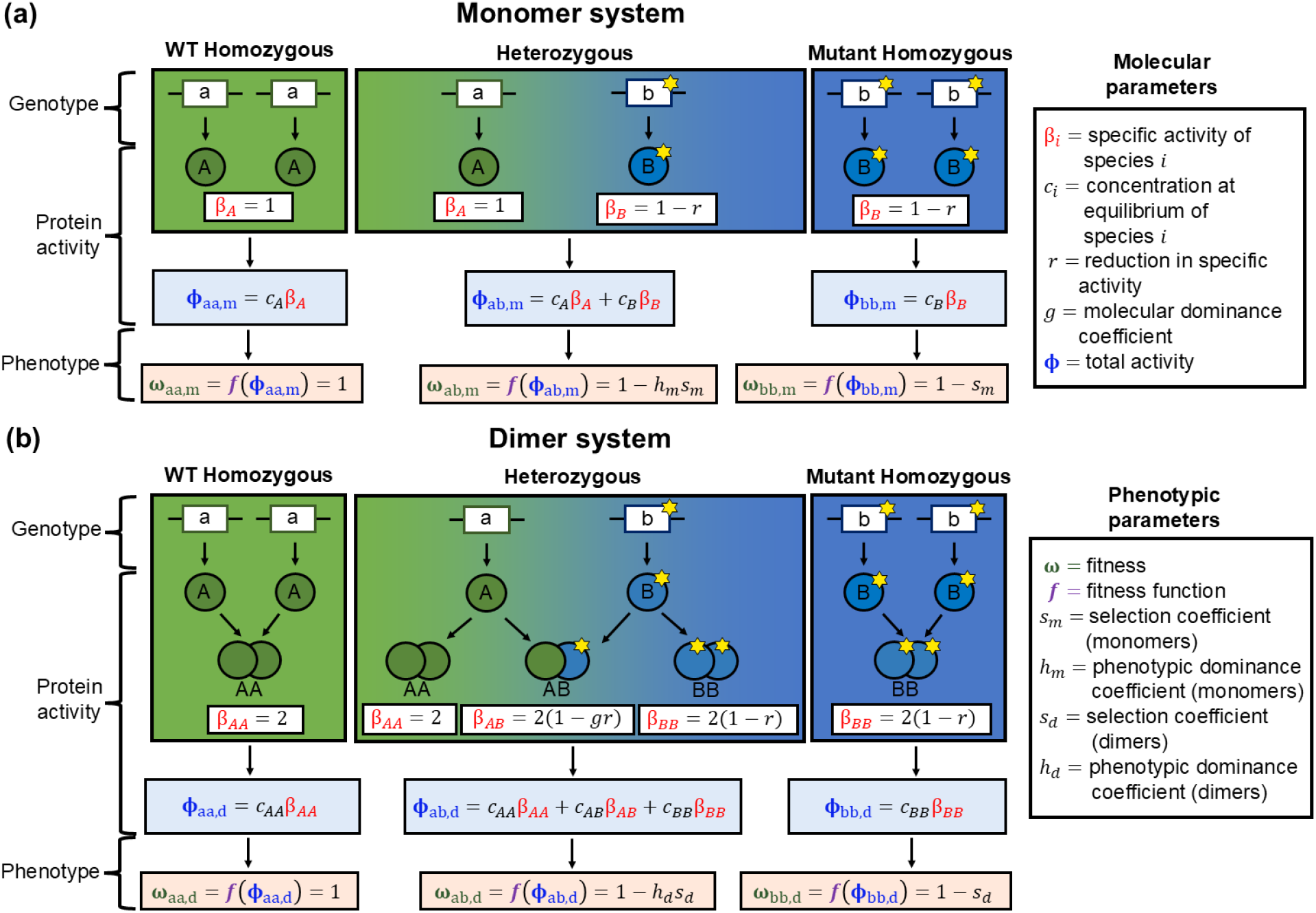
Effect of molecular dominance on phenotypic dominance. **a-b)** Dominance model at the molecular and phenotypic levels for genes encoding monomers (**a**) or dimers (**b**). In both cases, the model considers three possible genotypes with two copies of a focal gene: WT homozygote, heterozygote (WT + Mutant), and mutant homozygote. The specific activity of the WT monomer is set to 1, whereas the mutant monomer is set to 1 -*r*. Molecular dominance results from the physical interaction between the two proteins produced from the two alleles, thus modifying the specific activity of the heterodimer using the molecular dominance coefficient (*g*). The total protein activity is calculated as the weighted sum of the specific activity values of each functional unit (monomers in panel **a**, dimers in panel **b**), with their respective concentrations as weights. Fitness is then inferred based on a fitness function for the estimated values of total activity. Phenotypic dominance is observed in the two heterozygotes, with the value of the phenotypic dominance coefficient (*h*) being inferred based on the fitness values (phenotype) of the two respective homozygotes and the heterozygote.

In contrast, in the case of dimers, the two proteins form a mix of homo- and heterodimers (**Fig. 1b**). We assume that each subunit in the dimer contains an active site, so that dimers have twice the activity of a monomer. Based on a recent survey of structures in the Protein Data Bank, the ligand binding sites of around 80% of homomeric proteins involve only one subunit at a time (Abrusán and Marsh 2018), so most homomeric proteins meet our assumption. Thus, the specific activity of the mutant homodimer, *β*_*BB*_ is given by a modified version of equation 3:

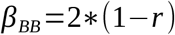

The dimer system introduces the possibility of heteromerization, so we define molecular dominance based on how similar the activity of the heterodimer is to that of the mutant homodimer, as shown in equation 4:

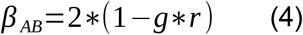

where *β*_*AB*_ indicates the activity of the heterodimer, and *g* indicates the molecular dominance of the mutant allele. This definition is analogous to that of phenotypic dominance and provides a comparison between different levels of effects of the mutant allele on the heterodimer. For example, a molecularly dominant loss-of-function allele that produces an inactive homodimer and inactivates the heterodimer would be described by *r* = 1 and *g* = 1. Conversely, a molecularly recessive loss-of-function allele that produces an inactive homodimer but that has no effect on the heterodimer would be described by *r* = 1 and *g* = 0.

In both systems, the total protein activity (ϕ). is calculated based on the total number of active proteins and their specific activity (see equation 5 in Methods). For simplicity, in our dimer systems, we assume that all protein copies assemble into stable homo- or heterodimers. Since the dimer system would also have half the concentration of dimers as that of monomers in the monomer system, our assumption of each dimer having two active sites allows comparing directly the total activities of both systems. Total activity can be linked to fitness (*ω*) with the given fitness function (*f*) of the system of interest (see equation 6 in Methods). However, since fitness functions can vary broadly and often independently from the effects of molecular interactions, we focus first on total activity.

In the rest of our study, we examine how mutational effects propagate through the different levels - molecular to phenotypic - in our model. In the heterozygous monomer system, total activity results from the net sum of the effects of the two monomers. Assuming an equal and constant number of copies of A and B, total activity becomes entirely dependent on the value of *r* (see equations 7 and 8 in Methods). In turn, the total activity in the heterozygous dimer system depends on the effects of the two homodimers and the heterodimer. Assuming identical binding affinities, free mixing, and an equal number of copies of A and B, the expected fractions at equilibrium would be 50% heterodimers and 25% of each of the homodimer (Reiner A. Veitia 2007; Hochberg et al. 2018; R. A. Veitia, Caburet, and Birchler 2018; Cisneros et al. 2024). Because of heterodimerization, the total activity in the dimer system becomes dependent on both *r* and *g* (see equations 9 and 10 in Methods). Ultimately, total activity is used to determine fitness using a fitness function. In the following sections, we explore how changes to *r, g*, and the proportions of the molecular complexes at equilibrium impact total activity. Finally, we show how fitness and phenotypic dominance emerge from *r, g*, and the corresponding fitness function.

## Analysis

### Molecular dominance changes the landscape of total protein activity

We explored the effect of physical interactions on total activity. As a reference, we first tested how changing values of *r* modify the total protein activity of homo- and heterozygotes encoding monomeric proteins. As noted above, equations 7 and 9 show how the selection coefficients for WT and mutant homozygotes in both systems are identical because they produce the same total activity (dashed black lines in **Fig. 2**). In the case of heterozygotes, the general effect of *r* on total activity in the dimer system is similar to that in the monomer system, but the magnitude of the slope depends on the value of *g* (**Fig. 2, Fig. S1a-b**). Thus, the relationship between total activity and *r* for heterozygotes in the monomer system is restricted to the dashed green line in **Fig. 2**, whereas in the dimer system the same value of *r* could correspond to different values of total activity depending on *g*. We refer to the difference between activities of monomers and dimers (ϕ). In the case of_dimer_ - ϕ). In the case of_monomer_) as the homodimerization effect (**Fig. S1c**), which can be expressed analytically as a function of *g* (see equation 11 in Methods). When *g* = 0.5, the dimer system becomes additive and identical to the monomeric heterozygote (ϕ). In the case of_dimer_ - ϕ). In the case of_monomer_ = 0) because the specific activity of heterodimers matches the average of those of the two homodimers. Since the dimer system is expected to form 50% heterodimers and 25% of each homodimer, its total activity becomes identical to that of the monomer system. On the other hand, mutational effects on total activity in the dimer system are potentiated when *g* > 0.5 and buffered when *g* < 0.5. Overall, our model shows how the possibility of molecular dominance can lead to different total activities from alleles that have the same specific activity.

**Figure 2.**
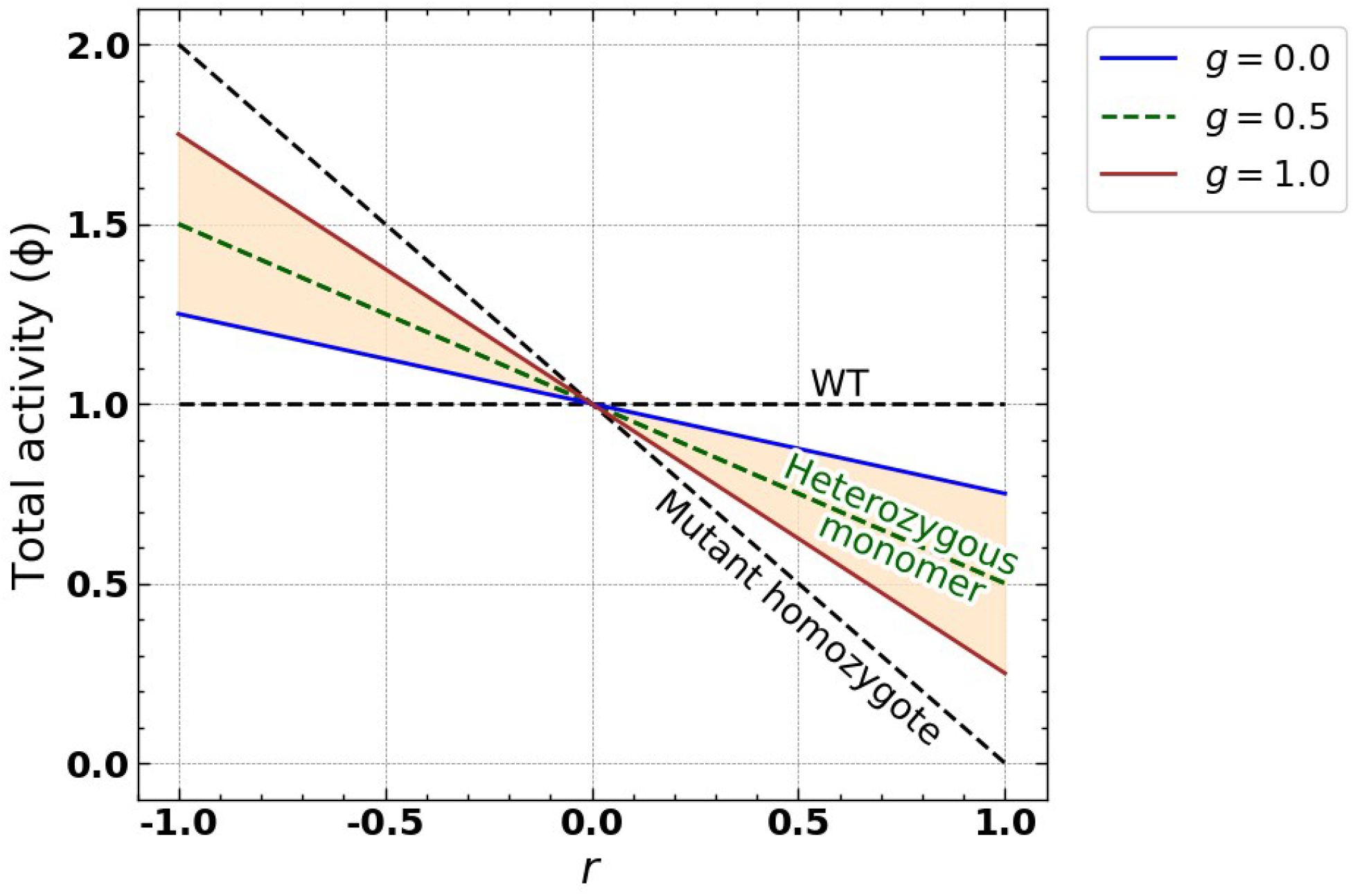
Comparison between the landscapes of total activity for the monomer and dimer systems. Total activity in the monomer and dimer systems as a function of *r*, which could change as a consequence of mutations. Dashed black lines indicate the total activity landscapes for WT and mutant homozygotes for both the monomer and dimer systems. In the monomer system, the heterozygote landscape is given by the dashed green line. The shaded region indicates the space accessible to heterozygotes in the dimer system depending on the value of molecular dominance (*g*), delimited by the blue (*g* = 0) and orange lines (*g* = 1).

Phenotypic dominance results from the heterozygote having the same fitness as one of the homozygotes. Since fitness is a function of activity, we sought to identify the values of molecular dominance that would produce identical activities in homo- and heterozygotes, which would then imply phenotypic dominance. For dimeric proteins, a molecularly dominant (*g* = 1) mutation implies the heterodimer has the same specific activity as the mutant homodimer, but it does not directly translate to the heterozygote having the same fitness as the mutant homozygote. This occurs because the heterozygous dimer system also contains a fraction of WT protein complexes, which contributes to the differing total activity (and hence fitness) of the heterozygote and mutant homozygote, even with the heterodimer having the same specific activity as the mutant homodimer (**Fig. 2**). Instead, the heterozygote and mutant homozygote systems become identical at the phenotypic level at *g* = 1.5 (**Supp. Video 1**), which would indicate a stronger change in specific activity for the heterodimer than for the mutant homodimer. When *g* = 1.5, the systems become identical because the sum of specific activities weighted by the relative concentrations of each dimer become identical to that of the mutant homozygote, regardless of the value of *r*. Conversely, heterozygotes in the dimer system have the same total activity as a WT homozygote when *g* = -0.5, also regardless of the value of *r* (**Supp. Video 1**). These results show that molecular dominance does not map 1:1 to phenotypic dominance due to the mix of homo- and heteromers at equilibrium.

The above cases imply that the effect of the mutation is potentiated (*g* > 1) or goes in the opposite direction (*g* < 0) in the heterodimer. While the extent to which such mutations occur in nature is unknown, there are observations that seem analogous to both cases. Some combinations of mutations in the homodimeric protein leucine aminopeptidase produce heterozygotes with higher catalytic activity than the two homozygotes (Sarver, Katoh, and Foltz 1992). Phenotypic overdominance is defined as the case in which the heterozygote has a higher fitness than the two homozygotes (Di and Lohmueller 2024; Manna, Martin, and Lenormand 2011), so extending our analogy molecular overdominance would be defined as the case when the heterodimer in a heterozygous dimer system has a higher catalytic activity than the two homodimers. The leucine aminopeptidase alleles would then be examples of observed phenotypic overdominance that is most likely driven by molecular overdominance, with *r* < 0 and *g* > 1.5. Overall, in our model, molecular overdominance occurs when *r* < 0 and *g* > 1, as well as when *r* > 0 and *g* < 0. Examples of negative values of *g* could be analogous to reports of loss-of-function mutations that complement each other by forming heteromers in enzymes like arginosuccinate lyase (Walker et al. 1997), type B dihydrofolate reductases (Dam et al. 2000), and cytosine deaminase (Després et al. 2024). Our model provides a framework to describe these cases, although they may be rare.

### Changes in equilibrium concentrations of homo- and heterodimers modify dominance effects

Having established the differences between homo- and heterozygotes, we focused on other factors specific to heterozygous alleles encoding dimers. The encoded proteins in this system are expected by default to form 50% heterodimers and 25% of each homodimer at equilibrium. Since mutations could modify this equilibrium by altering the binding affinities of homo- and heterodimers (Hochberg et al. 2018; Cisneros et al. 2024; Cortez-Romero et al. 2025), they could modify the impact of molecular dominance. If heteromerization is abolished, the two encoded proteins would become independent of each other like in our monomer system, effectively becoming additive and eliminating the effect of molecular dominance. In turn, if the proteins only heteromerize, then all dimers would be susceptible to the effects of mutations and molecular dominance.

We examined how shifts in the relative concentration of heterodimers influence total activity (**Fig. 3, Fig. S2**). For mutations with constant *r* but varying *g*, higher fractions of heterodimers result in slopes of a larger magnitude in the direction of *r* (**Fig. 3a-b**). All the lines intersect when *g* = 0.5, showing that the effect of higher heterodimer fractions disappears when the heterodimer activity is the average of the two homodimers. Indeed, the observed activity is the same as with 0% heterodimers, which is also the same as observed in the additive monomer model. Another interesting point in the curves is the value of *g* required to mask the effect of any mutation given a particular equilibrium of homo- and heterodimers. Since the concentration of both homodimers tends to remain identical to each other as long as both of them are stable (Cisneros et al. 2024; Cortez-Romero et al. 2025), the required values of *g* can be established analytically as a function of the relative fraction of heterodimers (equation 12 in Methods). Our model thus suggests that, given a specific combination of molecular dominance and heterospecificity, any mutation could produce the same activity as the WT homozygote and be phenotypically recessive. Similarly, we derived expressions for the selection and dominance coefficients for any combination of relative concentrations of homo- and heterodimers and any fitness function (see Supplementary Text). Overall, the effects of mutations on total activity become more pronounced for higher values of *g* and higher relative concentrations of heterodimers.

**Figure 3.**
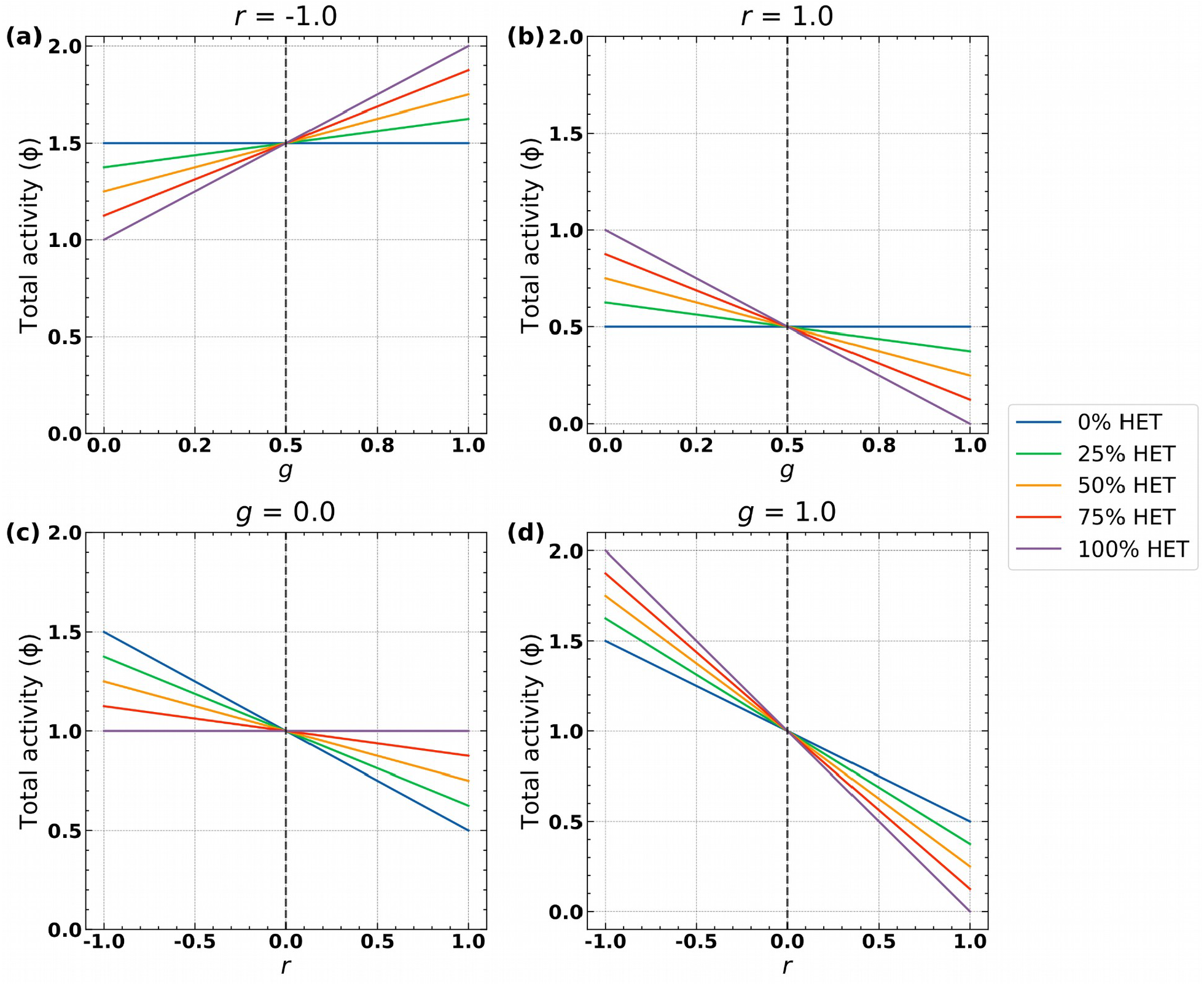
The equilibrium between homo- and heterodimers modulates the activity landscape. **a-b)** Total activity observed for heterozygotes harboring mutant alleles with varying molecular dominance values and effects on activity fixed to *r* = -0.5 (**a**) and *r* = 0.5 (**b**). **c-d)** Total activity observed for heterozygotes harboring mutant alleles with varying effects on activity and molecular dominance values fixed to *g* = 0 (**c**) and *g* = 1 (**d**).

The interaction between heterospecificity and molecular dominance yielded a different behavior. A higher relative heterospecificity buffered the effects of molecularly recessive mutations (*g* = 0, **Fig. 3c**) and potentiated those of molecularly dominant ones (*g* = 1, **Fig. 3d**). These inverse effects happen because heterodimers containing molecularly recessive mutations have a more similar activity to WT homodimers, whereas those harboring molecularly dominant mutations have a more similar activity to the mutant homodimers. Intuitively, these effects disappear when mutations have no effects on activity (*r* = 0), as shown by the intersection of all the lines.

Our model of the interactions between *r, g*, and the percentage of heteromers is consistent with several empirical observations. For example, proteins that selectively homomerize tend to be depleted in dominant negative mutations (Badonyi and Marsh 2023). In our model, the expected effects on total activity of such dominant negative mutations (*r* > 0, *g* = 1) become smaller as the fraction of heteromers decreases. Conversely, heterodimers of the paralogs RAF1 and BRAF1 promote MEK/ERK signalling more strongly than RAF1 homodimers. Mutations in RAF1 that enhance its heteromerization with BRAF1 then result in increased signalling (X. Wu et al. 2012). In our model, increased heteromerization also potentiates the effects of such mutations (*r* < 0, *g* > 0.5). Together, these results demonstrate how higher heterospecificity amplifies the effects of mutations with higher molecular dominance, while mitigating those with lower molecular dominance.

### Fitness and phenotypic dominance landscapes under different fitness functions

Since our ultimate goal is to connect dominance at the molecular level to dominance at the phenotypic level, we now connect molecular activity to phenotype, here specifically focusing on fitness. Multiple studies have shown that the relationship between protein activity and fitness is diverse, highly gene-specific (Keren et al. 2016) and environment-dependent (Dekel and Alon 2005; Després et al. 2022). Thus, we use the example of the cytosine deaminase protein (Fcy1) in two different growth conditions to show how fitness functions and molecular dominance contribute to phenotypic dominance. Briefly, Fcy1 catalyzes the conversion of cytosine into uracil, which allows the growth of auxotrophic yeast strains whose *de novo* uracil synthesis pathway has been disrupted. However, Fcy1 is also the target of antifungal drug 5-fluorocytosine (5-FC). Fcy1 catalyzes the conversion of 5-FC into 5-fluorouracil, which causes

DNA damage through thymine starvation and disrupts messenger RNA synthesis and protein translation (Longley, Harkin, and Johnston 2003). As a result, Fcy1 activity is toxic in media with 5-FC. We derived fitness functions by fitting a Hill equation to the data from Després et al., (2022) on yeast growth at different concentrations of cytosine and 5-FC, assuming they reflect the fitness function one would obtain by varying Fcy1 activity levels. We normalized the data on various concentrations of each molecule to represent equivalents of Fcy1 activity and assigned a WT reference fitness of 1 (**Fig. 4a-b**) (see full derivation in Methods).

**Figure 4.**
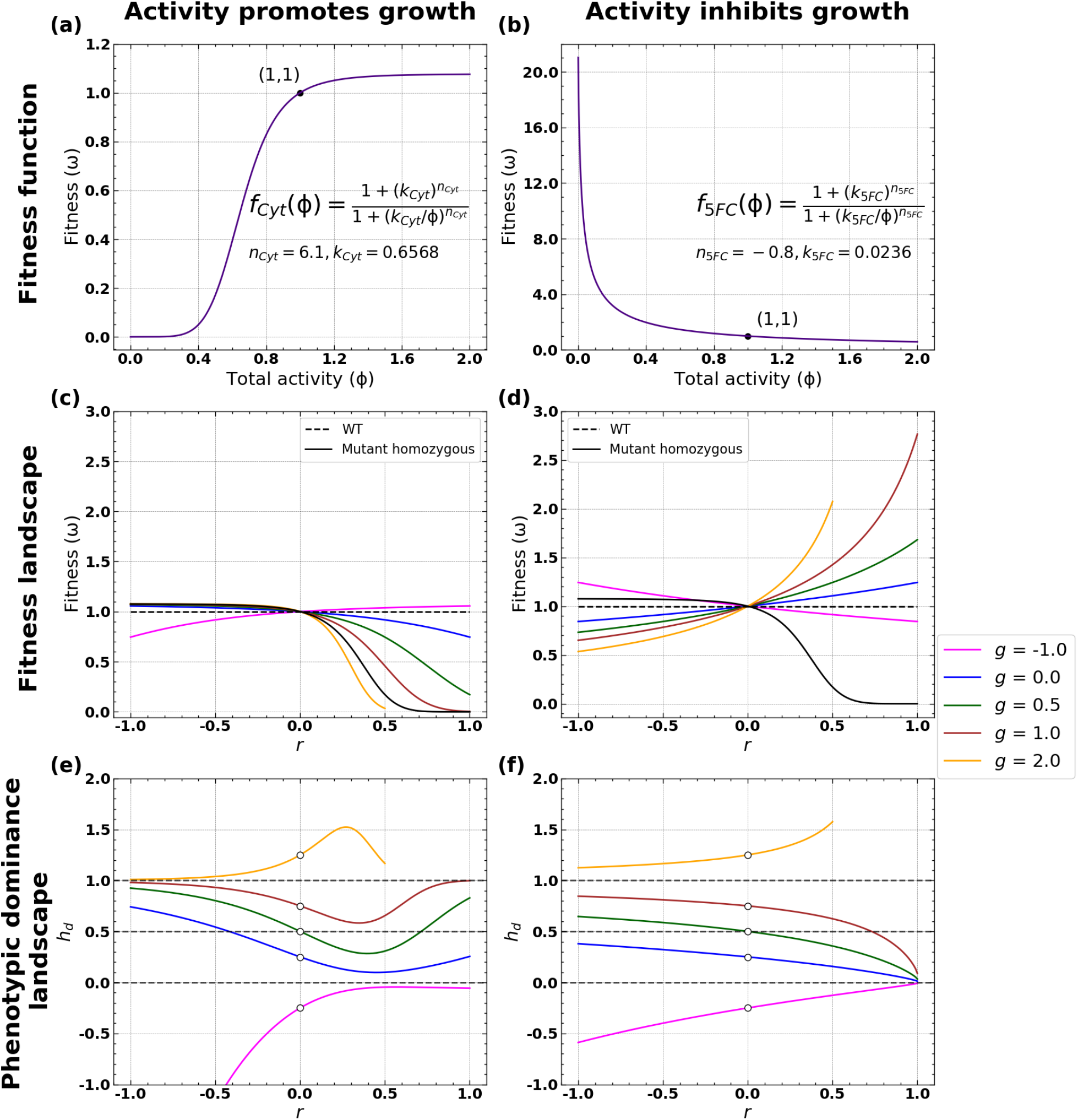
Effects of physical interactions on fitness and phenotypic dominance given two different fitness functions. **a-b)** Empirical fitness functions derived from Després et al., (2022) for yeast growth in media with cytosine (**a**) and media with 5-fluorocytosine (5-FC) (**b**). The activity of enzyme Fcy1 promotes growth in media with cytosine and inhibits growth in media with 5-FC. **c-d)** Fitness landscapes for mutations with different *r* and *g* values for each corresponding fitness function. The dashed horizontal line indicates a fitness of 1, which corresponds to the WT. **e-f)** Phenotypic dominance landscapes for mutations with different *r* and *g* values for each corresponding fitness function. Dashed horizontal lines indicate *h*=0 (phenotypic recessivity), *h*=0.5 (additivity), *h*=1 (phenotypic dominance). The fitness and phenotypic dominance curves for *g*=2.0 stop after *r*=0.5, as beyond that point, the heterodimer would have a negative specific activity, which is not allowed.

We examined how modifying the molecular parameters *r* and *g* affected fitness given each fitness function (**Fig. 4c-d, Fig. S3a-b**). Since different combinations of *r* and *g* values can provide the same activity value, they also have identical fitness effects. Similarly, as shown for the effects on protein activity in **Fig. 2**, different values for the molecular dominance parameters allow the system to reach fitness values that are inaccessible to the monomer system (equivalent to the case where *g* = 0.5 in **Fig. 4c-d**). In general, the effects of mutations on fitness are potentiated when *g* > 0.5 and buffered when *g* < 0.5. Our model also shows cases of phenotypic over- and underdominance, corresponding to instances in which the fitness of heterozygotes is respectively higher or lower than that of both the WT and the mutant homozygote (**Fig. 4c-d**). When higher activity promotes growth, underdominance is caused by mutations with *r* > 0 and *g* > 1.5, as well as mutations with *r* < 0 and *g* < -0.5 (**Fig. 4c**). In turn, overdominance is driven by mutations with *r* > 0 and *g* < -0.5, as well as mutations with *r* < 0 and *g* > 1.5 (**Fig. 4c**). When higher activity inhibits growth, these same mutations produce under- and overdominance but in the opposite direction (**Fig. 4d**).

We asked if there is an inherent relationship between molecular and phenotypic dominance. As defined above in equations 1 and 2, phenotypic dominance is calculated directly from observed differences in fitness (Greenberg and Crow 1960). We calculated the phenotypic dominance coefficients (*h*_*d*_) along the two fitness landscapes (**Fig. 4e-f, Fig. S3c-d**). Note that the phenotypic dominance coefficient is undetermined for mutations that have no effect (*r* = 0). As shown by the curves and as expected based on previous models (Kacser and Burns 1981; Manna, Martin, and Lenormand 2011; Xie et al. 2023), phenotypic dominance can arise directly from the fitness function without the need for molecular dominance arising from physical interactions (curve for *g* = 0.5). However, our models show that the relationship between molecular and phenotypic dominance is specific to each fitness function so one cannot easily relate one to the other without knowledge of the fitness function. For example, when using the fitness function for growth in cytosine (**Fig. 4e**), molecular dominance imposes a minimum value on the phenotypic dominance coefficient that is not observed in the curve for 5-FC (**Fig. 4f**). Using equations 8 and 10 that express the phenotypic dominance coefficient, we derived an expression that indicates the condition that the fitness function must satisfy to produce local minima or maxima (see Supplementary Text). These results show that while phenotypic dominance is influenced by molecular dominance, it is also shaped by properties of the underlying fitness function.

While on average higher molecular dominance tends to result in higher phenotypic dominance, we found some exceptions. Interestingly, in both fitness functions, some strong mutational effects would appear to invert the relationship between molecular and phenotypic dominance. For instance, a strong gain-of-function (*r* ∼ -1) with low molecular dominance (*g* = 0) in **Fig. 4e** can appear phenotypically dominant (*h* > 0.5). This behavior occurs due to the diminishing returns for higher activity in the corresponding fitness function (**Fig. 4a**). On the other hand, a strong loss-of-function mutation (*r* ∼ 1) with high molecular dominance (*g* = 1) in **Fig. 4f** can appear phenotypically recessive (*h* < 0.5) due to the high fitness penalty of the activity of any WT homodimers **Fig. 4b**, albeit less so than molecularly additive mutations (*g* = 0.5). Together, these observations show that interpreting observed phenotypic dominance as molecular dominance is not always correct, as it depends on the fitness function.

## Discussion

In humans and other diploid species, alleles at one locus combine to produce phenotypes. Multiple mechanisms have been proposed separately for the dominance relationships between alleles, including the non-linearity of the underlying fitness function (Wright 1934; Kacser and Burns 1981; Brown et al. 2009; Manna, Martin, and Lenormand 2011) and molecular interactions between encoded proteins (Bergendahl et al. 2019; Backwell and Marsh 2022; Herskowitz 1987; Zschocke, Byers, and Wilkie 2023). Under which conditions do we expect molecular interactions between two allelic proteins to translate into phenotypic dominance is not entirely known. It is thus difficult to infer how alleles may interact molecularly from phenotypic data on the inheritance of phenotypes. For instance, it might be tempting to connect observed phenotypic dominance to molecular dominance. However, we can only confidently confirm such an association by validating decreases in activity in heterozygotes and evaluating the fitness function.

Here, we developed a model that disentangles the contributions of physical interactions and fitness functions to phenotypic dominance by joining them into a single, quantitative framework. While non-linearities in the underlying biophysical properties of monomeric proteins and their fitness functions have been shown to produce complex patterns of phenotypic dominance (Xie et al. 2023), physical interactions open a broader range of possibilities. We show that molecular dominance allows biallelic loci encoding homomeric proteins to occupy regions of the activity landscape that would have been inaccessible had these loci encoded monomeric proteins. These effects of molecular dominance can be further potentiated if the two copies selectively heteromerize, which could happen if mutations increase the binding affinity of the heteromer relative to that of the homodimers (Cisneros et al. 2024; Cortez-Romero et al. 2025). Finally, using empirical fitness functions we show that molecular and phenotypic dominance are often correlated. However, the shape of the fitness function and the strength of mutational effects can cause deviations from this behavior.

Heteromerization is a key factor driving the impact of molecular dominance on fitness. We showed that molecularly dominant alleles that preferentially heteromerize with the WT have more pronounced effects on protein activity than those that do not. Since higher-order oligomers tend to form higher fractions of heteromers (Bergendahl et al. 2019; Backwell and Marsh 2022), they could be more susceptible to the effects of molecular dominance. Furthermore, the fitness and dominance effects of a given mutation could be modified by a prior mutation that impacts that equilibrium. Mutations altering the equilibrium could be both coding or regulatory. Different studies have shown that homo- or heterospecificity between homologous proteins can be achieved with very few mutations (Cortez-Romero et al. 2025; Emlaw et al. 2021; Ashenberg et al. 2011; Hochberg et al. 2018). In turn, when regulatory mutations cause one allele to be more highly expressed, its respective homodimer should increase in abundance (Cisneros et al. 2024). Other proteins could be less susceptible to changes in their assembly. For example, proteins that assemble co-translationally have been associated with isoform-(Bertolini et al. 2021), allele-(Badonyi and Marsh 2023) and paralog-specific (Mallik et al. 2025) homodimers.

As a result, such proteins do not interact with other alleles and are thus less likely to harbor dominant negative mutations (Badonyi and Marsh 2023). A final consideration is the possibility that only a fraction of protein copies assemble into dimers, with the remaining copies preserving some residual activity as monomers, which could occur for transient protein interactions (Nooren and Thornton 2003; Traut 1994). In this case, the contribution of dimers to total activity represented in our model would be scaled by the fraction of protein copies that do form dimers, which could buffer the effects of molecular dominance. Further research could focus on deconvoluting these factors and evaluating the extent to which they contribute to dominance.

Molecular dominance and fitness functions combine to shape the landscape of phenotypic dominance. While phenotypic dominance can result directly from the shape of the fitness function, in general higher magnitudes of molecular dominance result in stronger phenotypic dominance. Interestingly, our observation that fitness functions and molecular dominance impose minimum values on phenotypic dominance suggests that a mutation being phenotypically dominant depends on the focal gene. For example, it is possible that substitutions on homomeric proteins with multi-subunit binding sites have overall larger molecular dominance magnitudes than those with independent binding sites (Abrusán and Marsh 2018). However, our model also suggests that for sufficiently strong mutational effects, molecularly dominant mutations could be phenotypically recessive. We observe this unintuitive result in the case of the environment where higher protein activity leads to lower fitness (**Fig. 4f**) because the fitness function produces a steep penalty for a slight increase in activity. This does not occur in the other condition (**Fig. 4e**), which suggests that testing the same mutations in another environment with a different fitness function could modify the relationship between phenotypic and molecular dominance. Indeed, Xie et al., (2023) observed that a given mutation affecting binding affinity to a ligand could switch from phenotypically dominant to recessive at different ligand concentrations, which is analogous to studying environments with different fitness functions. Overall, linking molecular and phenotypic dominance requires knowledge of the fitness function.

Phenotypic over- and underdominance can also emerge in our model from the fitness function and molecular dominance. Since the total activity for the heterozygote respectively matches that of the WT and mutant homozygotes when *g*=-0.5 and *g*=1.5, then more extreme values of *g* should correspond to phenotypic over- and underdominance. This is the case for the empirical fitness functions we tested (Després et al. 2022), although whether each case corresponds to phenotypic over- or underdominance depends on the fitness function and the sign of *r*. Furthermore, while the empirical fitness functions used in our study are strictly increasing or decreasing (**Fig. 4a-b**), other more complex fitness functions (Dekel and Alon 2005; Keren et al. 2016) could produce different patterns of over- and underdominance. For example, consider a fitness function in which slight increases in activity over the WT could be beneficial until the point in which excess activity becomes costly, which could be the case for signalling pathways containing oncogenes (Nitulescu et al. 2018). In this case, a partially dominant mutation that increases activity would cause the heterozygote to have an intermediate activity value, corresponding to a higher fitness and thus overdominance. Our model provides a framework that could be used to describe cases of over- and underdominance and evaluate the contributions of molecular dominance and fitness functions to their occurrence in nature.

Current experimental methodologies provide simpler avenues to obtain high-throughput measurements of cellular growth rates than for specific protein activities. Since our model establishes the relationship between molecular dominance, fitness, and phenotypic dominance, we envision that it could help link measured fitness effects to underlying molecular effects. Such analyses could produce high-throughput maps of mutational effects given a fitness function and a small subset of validation experiments, ultimately providing a better understanding of how Mendelian genetics emerge from cellular biochemistry.

## Methods

### Phenotypic dominance as a function of molecular parameters

#### Definitions of molecular and phenotypic dominance

Our model (**Fig. 1**) seeks to illustrate how physical interactions can contribute to the emergence of dominance. To do so, we distinguish between two types of dominance: phenotypic and molecular. Phenotypic dominance is defined based on the differences in fitness of a mutant homozygote and a heterozygote relative to a WT homozygote (Greenberg and Crow 1960), as shown in equations 1-2:

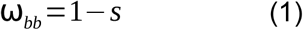

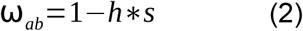

where *ω*_ab_ and *ω*_bb_ respectively indicate the fitness of the heterozygote and the mutant homozygote, *s* indicates the selection coefficient of the mutant homozygote, and *h* indicates the dominance coefficient of the mutant allele.

At the molecular level, we define the reduction in activity of the mutant monomer compared to the WT monomer (specific activity of 1) as *r*:

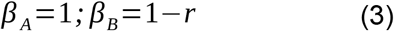

where *β*_*A*_ and *β*_*B*_ respectively indicate the specific activity of the WT and mutant monomer.

We assume activity to be additive at the protein level. Hence, the WT and mutant homodimers have twice the specific activity of the WT and mutant monomers, respectively. We define molecular dominance (*g*) as the effect of physical interactions between alleles on protein activity:

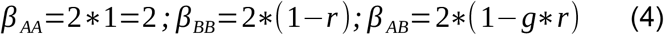

where *β*_*AA*_, *β*_*BB*_ and *β*_*AB*_ respectively indicate the activity of the WT homodimer, the mutant homodimer, and the heterodimer.

#### Calculation of total activity and fitness

For the system *j*, the total activity (ϕ). In the case of_j_) is calculated using the concentrations (*c*_*i*_) and the specific activities (*β*_*i*_) of each molecular species (*i*), as shown in equation 5:

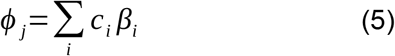

The fitness for each genotype (*ω*_j_) is then calculated based on the total activity using a fitness function (*f*) (**Fig. 4a-b**), as shown in equation 6 (see **Fitness function** section for more details):

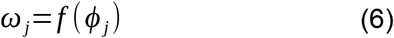

Selection (*s*) and phenotypic dominance coefficients (*h*) can then be estimated by comparing the fitness of the WT homozygote (*ω*_aa_ = 1) to that of the mutant homozygote (*ω*_bb_) and the heterozygote (*ω*_ab_), as shown in equations 1-2 (Greenberg and Crow 1960). These calculations of selection and phenotypic dominance coefficients apply to both the monomer (*s*_*m*_, *h*_*m*_) and dimer (*s*_*d*_, *h*_*d*_) systems.

#### Analytical expressions for the relationship between phenotypic and molecular dominance

We can use the above model to derive analytical expressions for the phenotypic parameters *s* and *h* in each system for any combination of molecular dominance (*g*), reduction in specific activity due to mutation (*r*), and fitness function (*f*):

##### WT homozygote, monomer system

Let *c* be the equilibrium concentration of the monomer A. The total activity (ϕ_aa,m_) would then correspond to *c*, since the specific activity of the WT monomers is 1. This represents the optimal activity of the WT homozygote, which we use as the normalization factor for all other systems. This is then used as input for the fitness function, which in return yields a unit fitness for the WT genotype. Thus, after normalizing by *c*, we have *f(*1*) =* 1.

##### Mutant homozygote, monomer system

Let *c* be the equilibrium concentration of the monomer B. Following equation 3, the specific activity of mutant monomers is 1 - *r*, which yields a total activity of *c*(1 - *r*). After normalizing by *c*, applying equation 6 and the fitness function we obtain equation 7, which directly expresses the selection coefficient as a function of the reduction in activity:

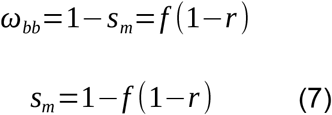

##### Heterozygous monomer system

Since now the two gene copies are different alleles, the concentration of each monomer will be *c*/2. Applying equation 5, the total protein activity becomes:

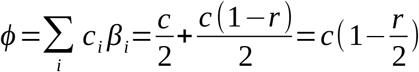

After normalizing by *c*, fitness becomes:

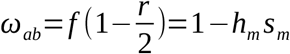

Since *s*_*m*_ is given by equation 7, the phenotypic dominance coefficient can be expressed as shown in equation 8:

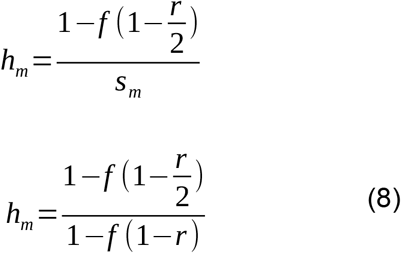

##### Homozygous dimer systems

Applying our above assumptions and assuming all protein copies assemble into dimers, the concentration of homodimers equals *c*/2. Since we allow dimers to have twice the specific activity of monomers to use the same normalization constant, we obtain that the total activity equals *c*. Thus, the homozygous dimer systems behave the same way as the monomeric systems: the fitness of the WT homozygote equals 1, and the selection coefficient of the mutant homozygote (*s*_*d*_) is calculated using equation 9:

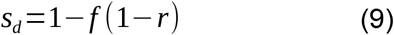

We notice here that the expression for the selection coefficient for homozygotes is identical in the monomer and dimer systems (equations 7 and 9).

##### Heterozygous dimer system

After applying our assumptions, the system is composed of *c*/2 total dimers at equilibrium. Since this is a 1:2:1 mixture of homo- and heterodimers, their concentrations are:

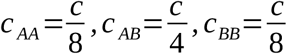

Substituting the specific activities from equations 3 and 4 into equation 5, the total protein activity is given by:

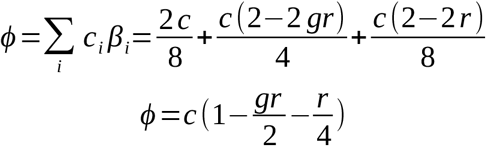

After normalizing by *c*, fitness becomes:

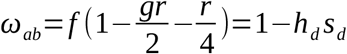

Since *s*_*d*_ is given by equation 9, the phenotypic dominance coefficient can be expressed as shown in equation 10:

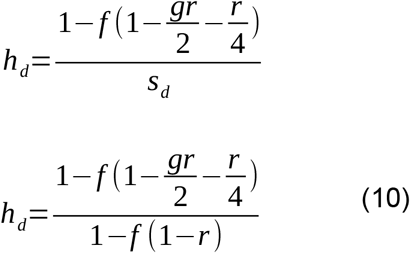

Equations 7-10 summarize how the selection and dominance coefficients can be obtained as a result of molecular dominance and the underlying fitness function.

### Homodimerization effect

We calculate the difference in total activity between the heterozygous dimer and monomer systems with identical reductions in activity and molecular dominance values. We refer to these differences in total activity between these otherwise identical systems as the homodimerization effect (Δϕ), which can be calculated analytically using equation 11: ϕ). In the case of), which can be calculated analytically using equation 11:

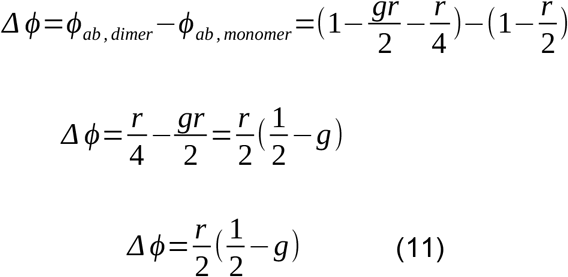

We set Δϕ), which can be calculated analytically using equation 11: ϕ). In the case of = 0 in the above equation to identify values of *r* and *g* that eliminate the homodimerization effect. We found two solutions: *r* = 0 and *g* = 0.5, For *r* = 0, the mutation has no effect on the activity of the protein, and thus we always obtain the same total activity regardless of the functional protein assembly. For *g* = 0.5, the heterodimer activity falls right in the middle of the WT and mutant homodimers, which results in a system equivalent to the heterozygous monomer, with half WT and half mutant species (**Supplementary Video 1**).

### Relationship between heterospecificity and masking of mutant alleles

A value of *g* that results in the masking of a mutant allele must provide the same total activity as the WT, which gives:

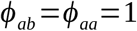

As shown in equation 5, the total activity of the heterozygous system is given by the sum of the contributions of all homo- and heterodimers. Assuming that *c* subunits assemble into *c*/2 dimers and that the two concentrations of homodimers are identical, we can establish the fraction of each homodimer in terms of the fraction of heterodimers:

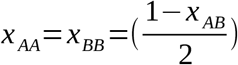

where *x*_*AA*_, *x*_*BB*_, and *x*_*AB*_ are the fractions of each of the two homodimers and the heterodimer, respectively.

Thus, the total activity of the heterozygous system can be written as shown in equation 12:

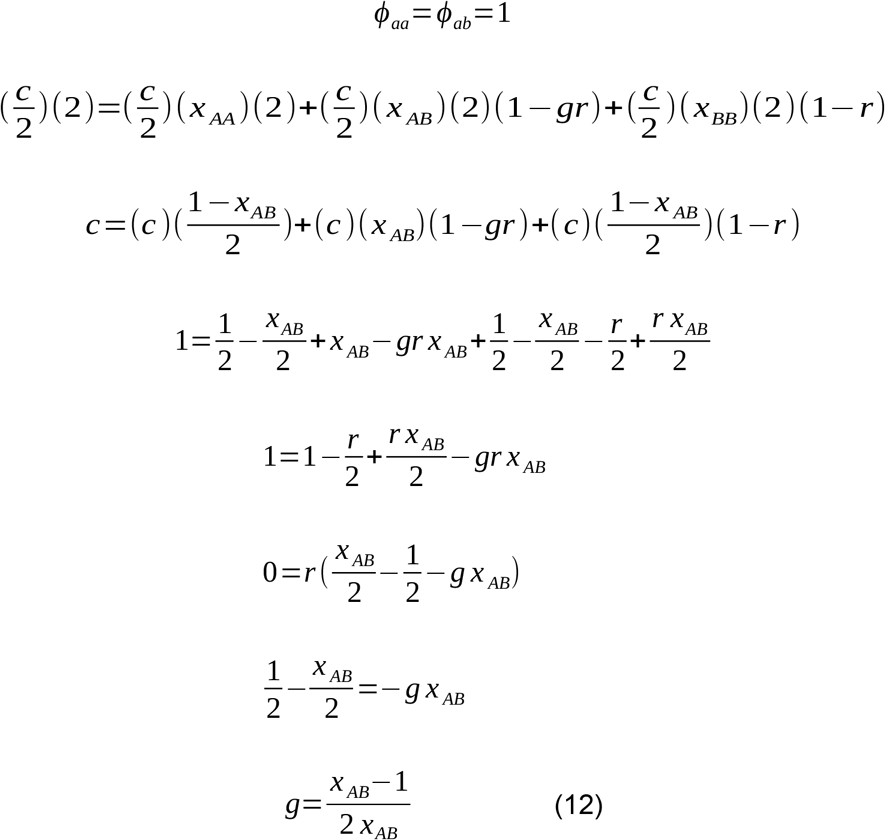

which indicates the value of *g* required to mask a mutant allele given the relative fraction of heterodimers. Importantly, this value is independent of the effect of the mutation on protein activity.

### Fitness functions

Fitness functions were derived from empirical data on cellular growth for yeast cells expressing Fcy1 presented in Després et al. (2022) (**Fig. 4a-b**). Briefly, yeast were grown with varying concentrations of one of two substrates of Fcy1: cytosine and 5-fluorocytosine (5-FC). Fcy1 catalyzes the conversion of cytosine into uracil, which in cytosine-rich media complements auxotrophic strains with disrupted *de novo* uracil synthesis pathways. As a result, Fcy1 activity becomes necessary for these strains to grow. Conversely, 5-FC is metabolized by Fcy1 into 5-fluorouracil, a toxic nucleotide analogue (Longley, Harkin, and Johnston 2003), so Fcy1 activity becomes toxic in media with 5-FC.

The data from Després et al. (2022) were fit to Hill equations, yielding the parameters:

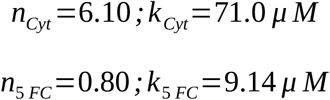

where *n* is the Hill coefficient, and *k* is the half-satuation constant. These parameters were obtained via fitting of the dose-response curves of relative growth rates (for cytosine medium) and relative inhibition rates (for 5-FC medium) against substrate concentration (in µM). The relative inhibition rates for 5-FC were subtracted from 1 to convert them into relative growth rates. The resulting curve fits a Hill equation with the same parameters, except the Hill coefficient is the additive inverse of its original value (*n*_*5FC*_ = -0.80).

Since higher concentrations of each substrate lead to higher protein activity, we reasoned that changes in protein specific activity with constant protein and substrate concentrations should follow similar fitness functions. Hence, we first scaled the substrate concentration data to represent equivalents of total protein activity. In our model, we deal with the range of *r* values from -1 to 1, and *g* values from 0 to 1. These values would represent the range of total activity from 0 (*r* = 1, loss-of-function mutation) to 2 (r = -1, a mutation that doubles protein activity), as seen in the mutant homozygous system (**Fig 2**). Thus, the substrate concentration values were scaled dividing by the maximum substrate concentration such that their range is 0 to 2 (in arbitrary units of total protein activity). The half saturation constants are also scaled similarly, to get the new values:

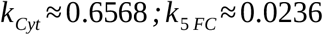

Then, we scaled the Hill-functions such that for a WT protein activity of 1, they return a WT fitness of 1 (ϕ). In the case of_aa_ = 1, *ω*_*aa*_ = 1) (**Fig. 4a**). This is done by equating the scaled function *f*(1)=1 and solving for the numerator (*A*):

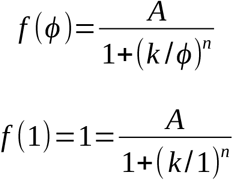

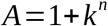

Thus, substituting the scaled parameter values, we get the fitness functions (*f*_*Cyt*_(ϕ). In the case of) and *f*_*5FC*_(ϕ). In the case of)), which are given by equations 13 and 14:

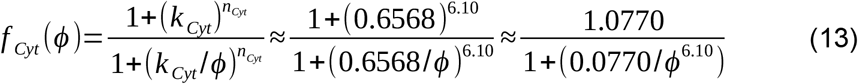

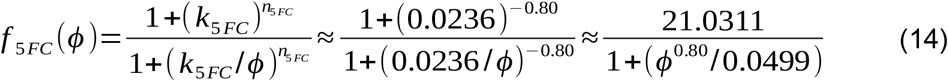

## Attributions

TP, AFC, and CRL designed research; TP, AFC, and CRL developed the theoretical model; TP, SD, and AFC analyzed data. TP, AFC, and CRL wrote the paper. CRL supervised the project.

## Acknowledgments

We thank the Landry lab members for their discussions during the project. We thank Simon Aubé, Pavithra Venkataraman, and Philippe Després for their comments on the manuscript.

## Data availability

All scripts used in this work are available at: https://github.com/Landrylab/Dominance_model_2025.git

## Supplementary Figures

**Figure S1.**
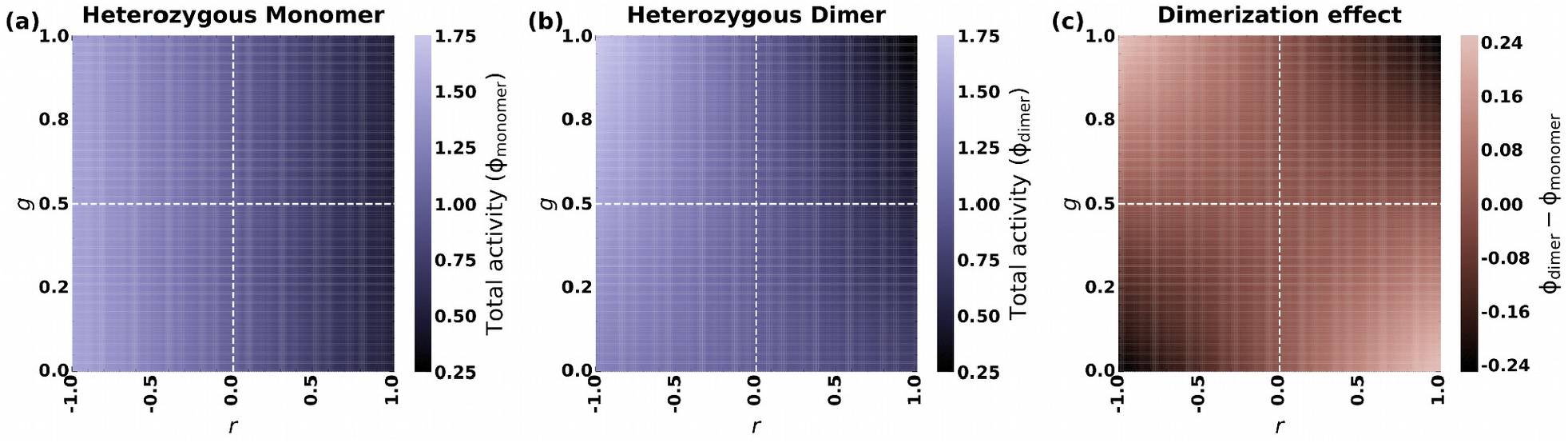
Extended comparison between the landscapes of total activity for the monomer and dimer systems. **a-b)** Activity landscapes of the heterozygous monomer (a) and dimer (b) systems as a function of *r* and *g*. Magnitude of the homodimerization effect (ϕ). In the case of_dimer_ - ϕ). In the case of_monomer_) as a function of *r* and *g*.

**Figure S2.**
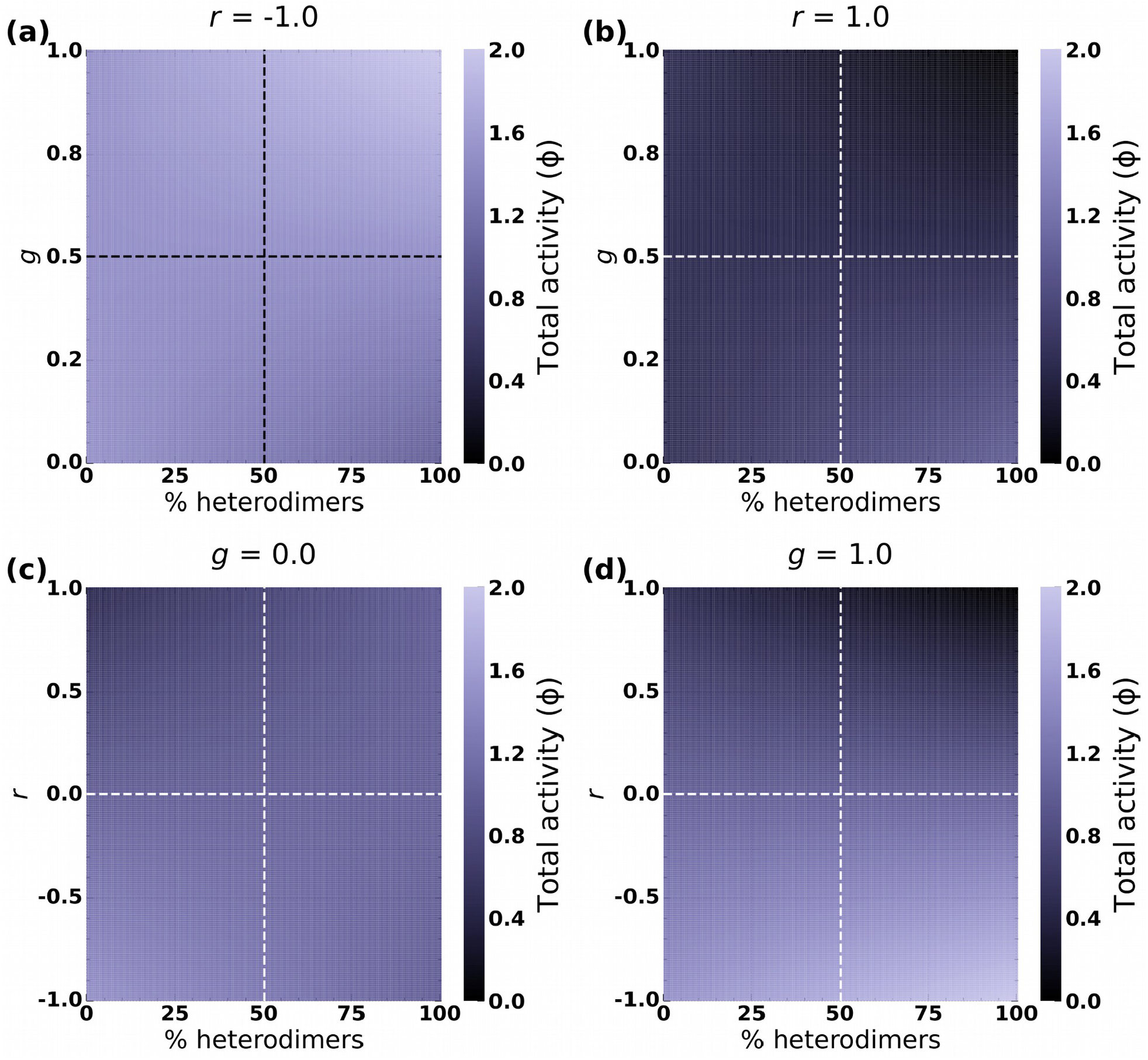
Extended view of the effect of the equilibrium of homo- and heterodimers on the activity landscape. **a-b)** Activity landscapes of the heterozygous dimer system as a function of the relative fraction of heterodimers and *g* when *r* = -1 (a) and when *r* = 1 (b). **c-d)** Activity landscapes of the heterozygous dimer system as a function of the relative fraction of heterodimers and *r* when *g* = 0 (c) and when *g* = 1 (d).

**Figure S3.**
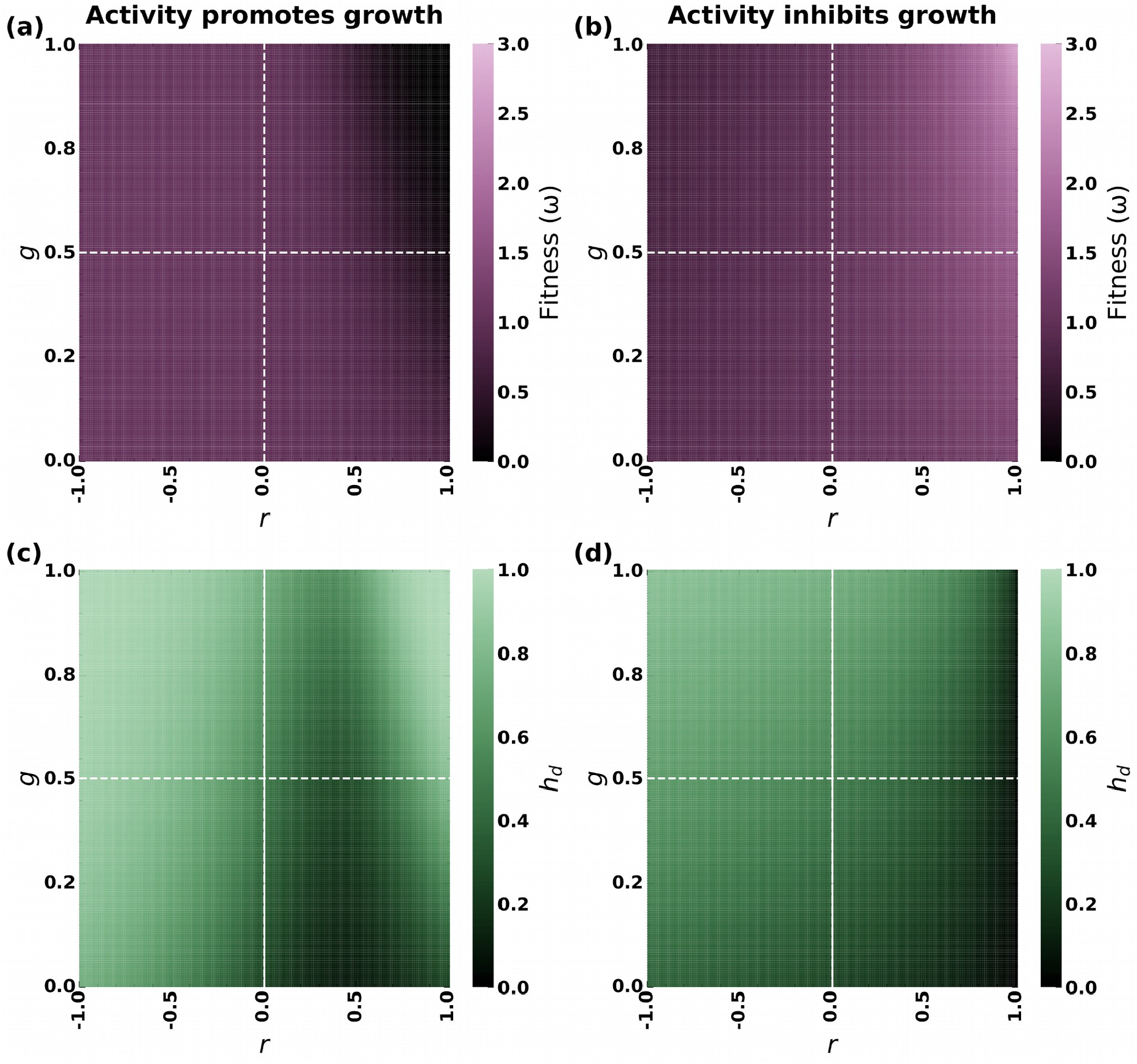
Extended view of the interaction between molecular dominance and different fitness functions. **a-** Fitness (a-b) and phenotypic dominance coefficient landscapes (c-d) of the heterozygous dimer system as a function of *r* and *g* using fitness functions derived from Després et al. (2022) and scaled to represent activity equivalents. As in Fig. 4, the fitness function for medium with cytosine was used in panels a and c, whereas the fitness function for medium with 5-FC was used in panels b and d.

**Supplementary Video 1.**
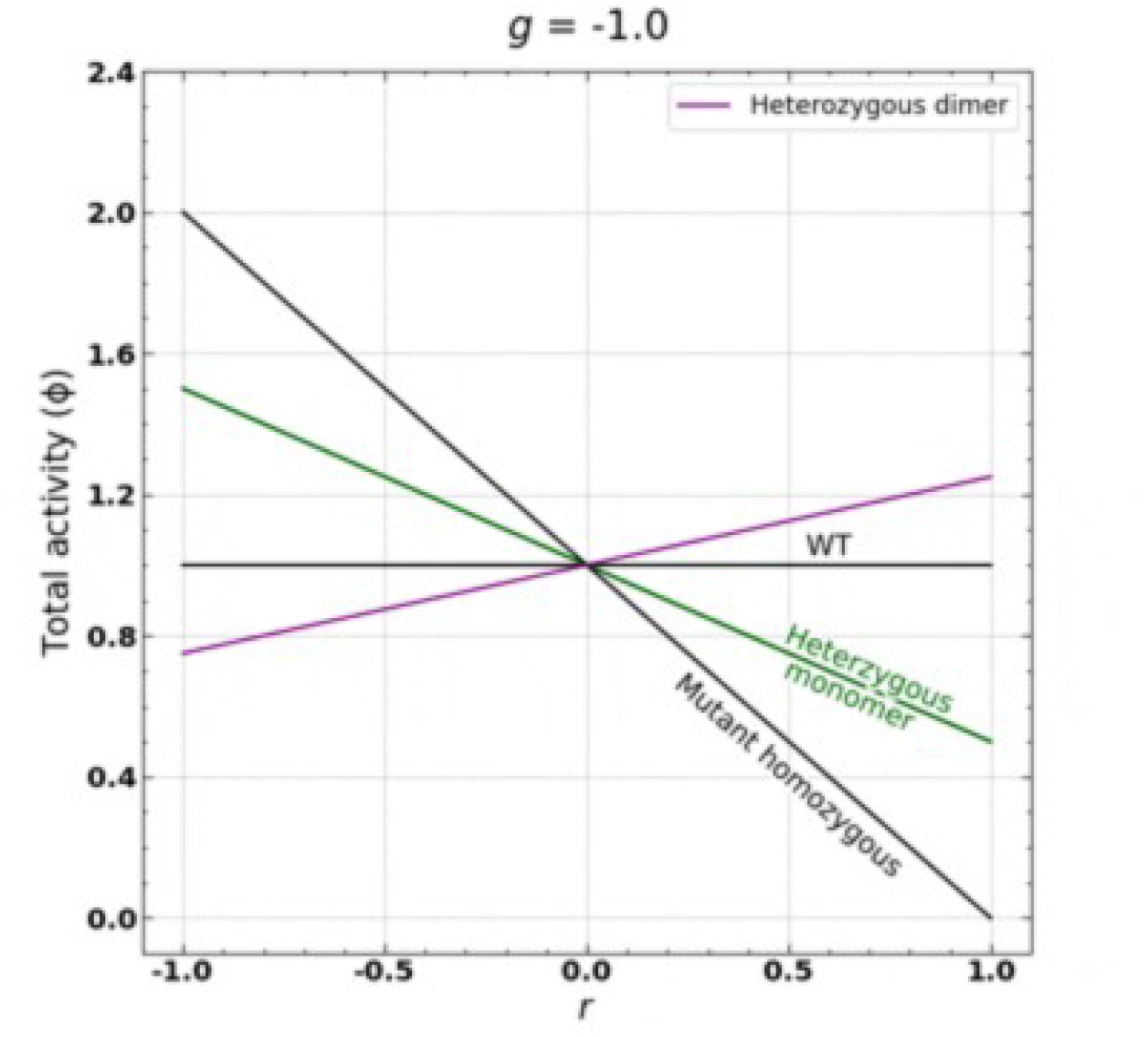
Comparison of total activity (ϕ) for the heterozygous dimer system under different values of *g* and *r* to other systems. While the activity of the homozygous mutant dimer (black) and the heterozygous monomer (green) systems are all determined solely by the reduction in specific activity (*r*), the heterozygous dimer system (purple) also depends on molecular dominance (*g*), which is varied from -1 to 2. The animation is paused briefly at particular values of *g* for which the total activity of the heterozygous dimer system is of interest: at ***g* = 0** (complete molecular recessiveness); at ***g* = 1** (complete molecular dominance); at ***g* = -0.5**, where it coincides with the WT phenotype, thus completely masking the effect of the mutant allele; at ***g* = 0.5**, it coincides with the heterozygous monomer phenotype, indicating the elimination of the homodimerization effect; at ***g* = 1.5**, it coincides with the mutant homozygous system, as the heterodimer’s effect on ϕ). In the case of overcompensates and nullifies that of the WT homodimer.

