## Supplementary_text for "Phenotypic dominance emerges from activity-fitness functions and molecular interactions"

### General expressions for phenotypic parameters as functions of molecular parameters

Let $k_a$ and $k_b$ respectively be the amount of protein product from a single allele.

*Monomer systems:*

1) WT homozygote:

Total protein activity is given by:

$$\phi_{aa,m} = c_A\beta_A = 2 * k_a * 1 = 2k_a$$

Since WT activity is set to 1 by definition, the normalizing factor for all systems becomes $2k_a$. Thus, normalized total activity becomes:

$$\phi_{aa,m} = \frac{2k_a}{2k_a} = 1$$

Fitness is given by:

$$\omega_{aa,m} = f(\phi_{aa,m}) = f(1) = 1 \text{ (by definition)}$$

2) Mutant homozygote:

Total activity:

$$\phi_{bb,m} = c_B\beta_B = 2 * k_b * (1 - r) = 2k_b(1 - r)$$

After normalizing:

$$\phi_{bb,m} = \frac{2k_b(1-r)}{2k_a} = \frac{k_b}{k_a}(1 - r)$$

Let $K$ be the amount of protein product from a mutant allele relative to a WT allele. Thus:

$$K = \frac{k_b}{k_a}$$

and normalized total activity becomes:

$$\phi_{bb,m} = K(1 - r)$$

Fitness:

$$\omega_{bb,m} = f(\phi_{bb,m}) = f(K(1 - r)) = 1 - s_m$$

Selection coefficient:

$$s_m = 1 - f(K(1 - r))$$

<u>3) Heterozygote:</u>

Total activity:

$$\phi_{ab,m} = c_A\beta_A + c_B\beta_B = k_a * 1 + k_b * (1 - r) = k_a + k_b(1 - r)$$

After normalizing:

$$\phi_{ab,m} = \frac{k_a + k_b(1-r)}{2k_a} = \frac{1+K(1-r)}{2}$$

Fitness:

$$\omega_{ab,m} = f(\phi_{ab,m}) = f(\frac{1+K(1-r)}{2}) = 1 - h_m * s_m$$

Phenotypic dominance coefficient:

$$h_m = \frac{1-f(\frac{1+K(1-r)}{2})}{s_m} = \frac{1-f(\frac{1+K(1-r)}{2})}{1-f(K(1-r))}$$

<u>*Dimer systems:*</u>

We assume complete dimerization for all the dimer systems, i.e., all the monomers assemble into dimers.

<u>1) WT homozygote:</u>

Total activity:

$$\phi_{aa,d} = c_{AA}\beta_{AA} = \frac{c_A}{2} * \beta_{AA} = \frac{2*k_a}{2} * 2 = 2k_a$$

After normalizing:

$$\phi_{aa,d} = \frac{2k_a}{2k_a} = 1$$

Fitness:

$$\omega_{aa,d} = f(\phi_{aa,d}) = f(1) = 1$$

<u>2) Mutant homozygote:</u>

Total activity:

$$\phi_{bb,d} = c_{BB}\beta_{BB} = \frac{c_B}{2} * 2 * (1 - r) = 2k_b(1 - r)$$

After normalizing:

$$\phi_{bb,d} = \frac{2k_b(1-r)}{2k_a} = \frac{k_b}{k_a}(1 - r) = K(1 - r)$$

Fitness:

$$\omega_{bb,d} = f(\phi_{bb,d}) = f(K(1 - r)) = 1 - s_d$$

Selection coefficient:

$$s_d = 1 - f(K(1 - r))$$

<u>3) Heterozygote:</u>

Here, $c_A = k_a$, $c_B = k_b$. Thus, total number of monomers = $k_a + k_b$, and the total number of dimers is half of that, i.e., $(k_a + k_b) / 2$. Let $x_{AA}$, $x_{AB}$, $x_{BB}$ be the percentages of WT homodimer, heterodimer, mutant homodimer respectively. Thus:

$$x_{AA} = \frac{c_{AA}}{(k_a+k_b)/2} = \frac{2c_{AA}}{(k_a+k_b)} \; ; \; x_{AB} = \frac{2c_{AB}}{(k_a+k_b)} \; ; \; x_{BB} = \frac{2c_{BB}}{(k_a+k_b)}$$

Thus, the equilibrium concentrations of the dimers expressed in terms of their percentages are:

$$c_{AA} = \frac{(k_a+k_b)}{2} * x_{AA} \; ; \; c_{AB} = \frac{(k_a+k_b)}{2} * x_{AB} \; ; \; c_{BB} = \frac{(k_a+k_b)}{2} * x_{BB}$$

Total protein activity:

$$\phi_{ab,d} = c_{AA}\beta_{AA} + c_{AB}\beta_{AB} + c_{BB}\beta_{BB}$$

$$\phi_{ab,d} = \frac{(k_a+k_b)}{2} * x_{AA} * 2 + \frac{(k_a+k_b)}{2} * x_{AB} * 2 * (1 - gr) + \frac{(k_a+k_b)}{2} * x_{BB} * 2 * (1 - r)$$

$$\phi_{ab,d} = \frac{(k_a+k_b)}{2} * 2 * \{x_{AA} + x_{AB} * (1 - gr) + x_{BB} * (1 - r)\}$$

$$\phi_{ab,d} = (k_a + k_b) * \{x_{AA} + x_{AB} * (1 - gr) + x_{BB} * (1 - r)\}$$

After normalizing:

$$\phi_{ab,d} = \frac{(k_a+k_b)}{2k_a} * \{x_{AA} + x_{AB} * (1 - gr) + x_{BB} * (1 - r)\}$$

$$\phi_{ab,d} = \frac{(1+K)}{2} * \{x_{AA} + x_{AB} - x_{AB} * gr + x_{BB} - x_{BB} * r)\}$$

$$\phi_{ab,d} = \frac{(1+K)}{2} * \{(x_{AA} + x_{AB} + x_{BB}) - x_{AB} * gr - x_{BB} * r)\}$$

$$\phi_{ab,d} = \frac{(1+K)}{2} * \{1 - x_{AB} * gr - x_{BB} * r)\}$$

Fitness:

$$\omega_{ab,d} = f(\phi_{ab,d}) = f(\frac{(1+K)}{2}\{1 - x_{AB}gr - x_{BB}r)\}) = 1 - h_d * s_d$$

Phenotypic dominance coefficient:

$$h_d = \frac{1-f(\frac{(1+K)}{2}\{1-x_{AB}gr-x_{BB}r)\})}{s_d} = \frac{1-f(\frac{(1+K)}{2}\{1-x_{AB}gr-x_{BB}r)\})}{1-f(K(1-r))}$$

### Fitness functions can impose minima or maxima on the phenotypic dominance coefficient

*Monomer system:*

Following equation 6, let *f*(ϕ) be a function that determines fitness (ω) with respect to total protein activity (ϕ). Similarly, following equation 8, let the dominance coefficient ($h_m$) be determined by the following expression:

$$h_m(r) = \frac{1 - f(1-\frac{r}{2})}{1 - f(1-r)}$$

where *r* is the effect of a given mutation on the specific activity of a monomer.

In order to look for a critical value $r_0$ such that $h'(r_0) = 0$, let $k(r) = ln(h_m(r))$. Thus:

$$h_m(r) = e^{k(r)}$$

$$h'_m(r) = e^{k(r)}k'(r)$$

Since $e^{k(r)}$ has no roots, $h'_m(r_0) = 0$ if and only if $k'(r_0) = 0$. Therefore, we look for a value $r_0$ such that $k'(r_0) = 0$.

Substituting the expression for $h_m(r)$ into $k(r)$, we obtain the following:

$$k(r) = ln(\frac{1-f(1-\frac{r}{2})}{1-f(1-r)}) = ln(1 - f(1 - \frac{r}{2})) - ln(1 - f(1 - r))$$

Using the above expression to calculate the derivative of $k(r)$:

$$k'(r) = \frac{1}{1-f(1-\frac{r}{2})}(- f'(1 - \frac{r}{2}))(- \frac{1}{2}) - (\frac{1}{1-f(1-r)})(- f'(1 - r))(- 1)$$

After algebraic manipulation:

$$k'(r) = \frac{f'(1-\frac{r}{2})}{2(1-f(1-\frac{r}{2}))} - \frac{f'(1-r)}{1-f(1-r)}$$

Equating to zero:

$$k'(r_0) = 0 = \frac{f'(1-\frac{r_0}{2})}{2(1-f(1-\frac{r_0}{2}))} - \frac{f'(1-r_0)}{1-f(1-r_0)}$$

$$\frac{f'(1-r_0)}{1-f(1-r_0)} = \frac{f'(1-\frac{r_0}{2})}{2(1-f(1-\frac{r_0}{2}))}$$

$$\frac{f'(1-r_0)}{f'(1-\frac{r_0}{2})} = \frac{1-f(1-r_0)}{2(1-f(1-\frac{r_0}{2}))}$$

The above condition must be satisfied for a given function to impose a local minimum or maximum on the phenotypic dominance coefficient in the monomer system. Additionally, $r_0$ must not be an inflection point for $k(r)$ and the denominators must be different from zero.

*Dimer system:*

We can apply a similar procedure to the equations for the dimer system. From equation 10, the phenotypic dominance coefficient is given by:

$$h_d = \frac{1 - f(1 - \frac{gr}{2} - \frac{r}{4})}{1 - f(1-r)}$$

where $r$ is the effect of a given mutation on the specific activity of a dimer and $g$ is the molecular dominance coefficient. As above, let $k(r) = ln(h_d(r))$. Thus:

$$h_d(r) = e^{k(r)}$$

$$h'_d(r) = e^{k(r)}k'(r)$$

Therefore, $h'_d(r_0) = 0$ if and only if $k'(r_0) = 0$. We then look for a value $r_0$ such that $k'(r_0) = 0$.

Substituting the expression for $h_d(r)$ into $k(r)$, we obtain the following:

$$k(r) = \frac{1 - f(1 - \frac{gr}{2} - \frac{r}{4})}{1 - f(1-r)} = ln(1 - f(1 - \frac{gr}{2} - \frac{r}{4})) - ln(1 - f(1 - r))$$

Using the above expression to calculate the derivative of $k(r)$:

$$k'(r) = \frac{1}{1-f(1-\frac{gr}{2}-\frac{r}{4})}(- f'(1 - \frac{gr}{2} - \frac{r}{4}))(- \frac{g}{2} - \frac{1}{4}) - \frac{1}{1-f(1-r)}(- f'(1 - r))(- 1)$$

After algebraic manipulation:

$$k'(r) = \frac{f'(1-\frac{gr}{2}-\frac{r}{4})}{1-f(1-\frac{gr}{2}-\frac{r}{4})}(\frac{g}{2} + \frac{1}{4}) - \frac{f'(1-r)}{1-f(1-r)}$$

Equating to zero:

$$k'(r_0) = 0 = \frac{f'(1-\frac{gr_0}{2}-\frac{r_0}{4})}{1-f(1-\frac{gr_0}{2}-\frac{r_0}{4})}(\frac{g}{2} + \frac{1}{4}) - \frac{f'(1-r_0)}{1-f(1-r_0)}$$

$$\frac{f'(1-r_0)}{1-f(1-r_0)} = \frac{f'(1-\frac{gr_0}{2}-\frac{r_0}{4})}{1-f(1-\frac{gr_0}{2}-\frac{r_0}{4})}(\frac{g}{2}+\frac{1}{4})$$

$$\frac{f'(1-r_0)}{f'(1-\frac{gr_0}{2}-\frac{r_0}{4})} = \frac{1-f(1-r_0)}{1-f(1-\frac{gr_0}{2}-\frac{r_0}{4})}(\frac{g}{2}+\frac{1}{4})$$

The above condition must be satisfied for a given function to impose a local minimum or maximum on the phenotypic dominance coefficient in the dimer system. Additionally, $r_0$ must not be an inflection point for $k(r)$ and the denominators must be different from zero.
